# The gap between elite female and male runners continues to shrink in the 21st century

**DOI:** 10.1101/2025.08.01.668047

**Authors:** Tommaso Jucker

**Author notes:** **Data and code accessibility statement** All data and R code underpinning the results of this study are publicly available on Zenodo: https://doi.org/10.5281/zenodo.18020910.

## Abstract

In 1992 a controversial analysis of historical trends in world record performances raised the question of whether women would one day run faster than men across a whole range of athletic disciplines. Since then, however, the consensus has been that the gap between male and female runners stabilised towards the end of the 20^th^ century and has since remained unchanged. Here I analyse running performances by elite athletes since 2001 to show that contrary to this widely held belief, women have in fact continued to close the gap on men across all distances from 60 m to the marathon. Women made the biggest gains relative to men in middle and long-distance running disciplines, and this occurred despite the fact that men too have continued to improve significantly in these events over the past 25 years. The jury is still out there as to whether women will one day run faster than men, but what does appear more certain is that women will continue to push the boundaries of running performance over the coming decades.

## Introduction

The limits of human running have long fascinated physiologists, biomechanical engineers, evolutionary biologists, statisticians and sports fans alike. In the 1990s and early 2000s, analyses of changes in running times by men and women during the 20^th^ century led to the bold prediction that women would outrun men in the not-too-distant future (1, 2). These controversial studies were heavily scrutinised (3, 4), with others arguing that this gap in running performance would never be bridged due to aerobic, biomechanic, musculoskeletal and hormonal differences between men and women (3–6). Since then, the general consensus has been that the gap between elite male and female runners is around 10% (3, 7, 8) and has stabilised around this value since the 1980s when women’s running professionalised and access to the sport increased (6, 9–11).

Here I question this assumption using a comprehensive dataset of 13,094 running performances recorded between 2001 and 2025 by elite male and female athletes across running events ranging in distance from 60 m sprints to the marathon. I show that contrary to conventional wisdom, the gap in athletic performance has shrunk markedly since the turn of this century, as female athletes have continued to improve faster than men across all running disciplines.

## Methods

All data formatting, analysis and visualisation were conducted in R (version 4.2.2) using the *rvest, xml2, tidyverse, lme4, lmerTest, plotrix* and *shape* packages. Data and R code to reproduce the results of this study are publicly available on Zenodo: https://doi.org/10.5281/zenodo.18020910.

### Athletic performance data

Results of all officially ratified races for running events that took place between 2001 and 2025 were downloaded from the World Athletics website (https://worldathletics.org) on the 21 December 2025. For the purposes of this analysis, I included 11 running events ranging in distance from 60 m to the marathon, which I categorized into sprint races (60 m, 100 m, 200 m and 400 m), middle-distance races (800 m, 1500 m and 3000 m), and long-distance races run on track (5 km and 10 km) and road surfaces (half marathon and marathon). Except for the 60 m, which are run indoors, for all other running distances I only included results from outdoor races. I excluded the mile, which is run less frequently and not contested at the Olympics or World Championships, as well any running events that include hurdles or barriers (as these events vary in distance and hurdle height between men and women). I also excluded all races run in 2020, as most were cancelled due to the Covid-19 pandemic (12).

For each running event, I first compiled the top 100 performances recorded in each year for both men and women. For short and middle-distance races, which are run most frequently, it is not uncommon for a handful of athletes to record many of the top performances in any given season. To avoid a small number of athletes having an undue influence on the trends, for each year I therefore only retained an athlete’s top performance in any given event. To ensure a similar number of performances were compared across running events, for each year I then subset the data to keep the top 25 performing athletes in each event for both men and women. This resulted in a total of 13,094 running performance recorded across 971 sporting events by 3611 elite athletes. Note that modifying these selection criteria to either retain a smaller (top 10) or larger (top 50) number of athletes per event in each year did not alter the results (Figs S1–2).

### Data analysis

For each year between 2001 and 2025, I calculate the mean time (in seconds) that the top 25 elite male and female athletes took to complete each running event. I then used these mean values to quantify the gap between men and women for each event and each year, both in terms of differences in pace (expressed in s km^-1^) and as a percentage.

To determine if differences in athletic performance between men and women are similar across running disciplines, I pooled all data from 2001 and 2025 and used ANOVAs to compare performance gaps among the 11 running events considered in this analysis.

To then test if differences in athletic performance between men and women have remained stable over the past 25 years, I used linear mixed effects models to characterise changes in performance gaps over time for each running event. Specifically, I modelled the performance gap between men and women as a function of calendar year, and allowed this relationship to vary among disciplines by including running event as a random intercept and slope term in the models (factor with 11 levels).

Finally, I used a similar mixed effects model to test how the mean running pace of elite men and women has changed between 2001 and 2025. This allowed me to determine if changes in the performance gap over time have arisen due to women making greater gains than men, or simply because men have stopped improving or slowed down in recent decades. As before, I allowed the relationship between running pace and calendar year to vary among disciplines by including running event as a random intercept and slope term in the models. Prior to model fitting, I visually inspected the data to rule out deviation from linearity and confirm that results were not affected by anomalous years in the timeseries that altered the strength and direction of trends (see Results for details).

## Results

The data show that the gap between elite male and female runners has shrunk significantly during the 21^st^ century across all running distances (Fig. 1 and Table S1). When averaged over the past 25 years, elite women’s running times were 12.6 % slower than men’s across disciplines. The size of the performance gap varied considerably among disciplines depending on their distance (Fig. 1 a–b), increasing progressively from short-distance sprint races such as the 60 m (9.2 %) until it reached a peak in middle and long-distance events such as the 10 km (14.2 %).

**Fig. 1:**
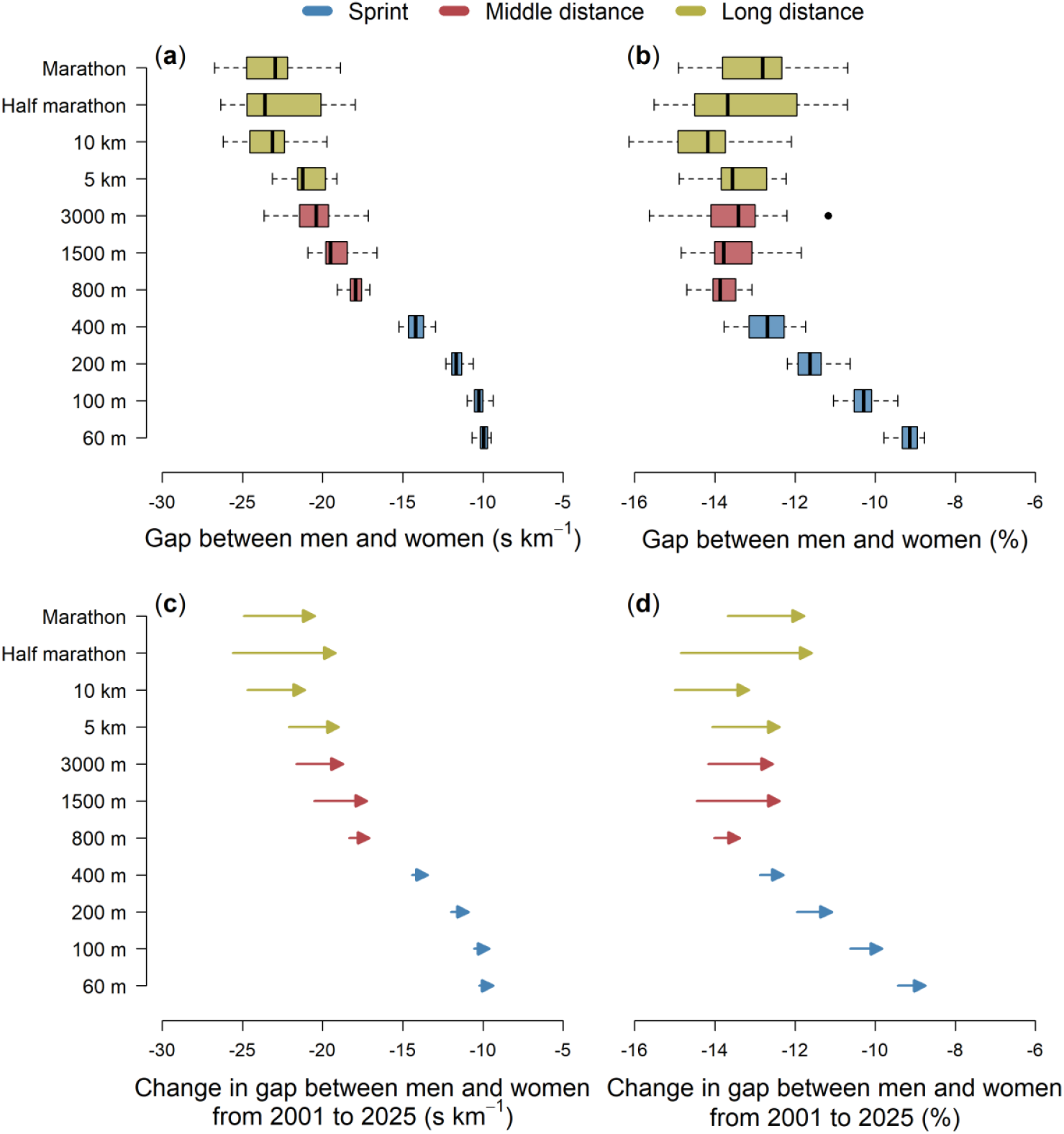
Shrinking gap among elite male and female runners during the 21^st^ century. Top panels depict variation in the gap – expressed as both a difference in pace between men and women (**a**) and as a % (**b**) – across the 11 events considered in this analysis for the period between 2001 and 2025. The more negative the values, the greater the gap between male and female runners. Bottom panels (**c**–**d**) show changes in the gap between 2001 (start of the arrow) and 2025 (arrow head) for each running event, as estimated from linear mixed effects models. Arrows pointing to the right indicate a shrinking gap over time, with the length of the arrow reflecting the magnitude of this change. Running events are colour-coded into sprint (blue), middle-distance (red) and long-distance races (yellow). Data underpinning estimated changes in the gap between male and female elite runners from 2001 to 2025 (**c**–**d**) are shown in Fig. 2.

Across all events, women significantly reduced this performance gap by an average of 1.3 % over the past 25 years – from 13.2 % in 2001 to 11.9 % in 2025 (Fig. 1 c–d). This shrinking gap between elite male and female runners was particularly pronounced in running events longer than 800 m, where women gained an average of 3.6 s km^-1^ against men’s times between 2001 and 2025 (equivalent to a 1.9 % reduction in the performance gap between elite male and female runners across these middle and long-distance events). As a result, the range and variability in the performance gap across running disciplines has visibly decreased since the start of the century, from 5.6 % in 2001 (maximum gap = 15.0 % in the 10 km; minimum gap = 9.4 % in the 60 m; standard deviation = 1.8 %) to 4.6 % in 2025 (maximum gap = 13.5 % in the 800 m; minimum gap = 8.9 % in the 60 m; standard deviation = 1.4 %).

Women reduced the gap with men even though elite male athletes have also continued to significantly improve their performances across all disciplines (Figs 2–3). Since 2001, male athletes have increased their running pace by an average of 2.2 s km^-1^ across disciplines, with the greatest gains coming in the half marathon and marathon (6.0–6.2 s km^-1^). But over the same period, women increased their average running pace by twice as much (4.5 s km^-1^ across disciplines, 10.3–12.6 s km^-1^ in the half marathon and marathon), making gains relative to men in all running events (Fig. 3). For both men and women, across almost all disciplines improvements in running performance were continuous and sustained over time, with no evidence of either inflection points or levelling off in the timeseries (Fig. 2).

**Fig. 2:**
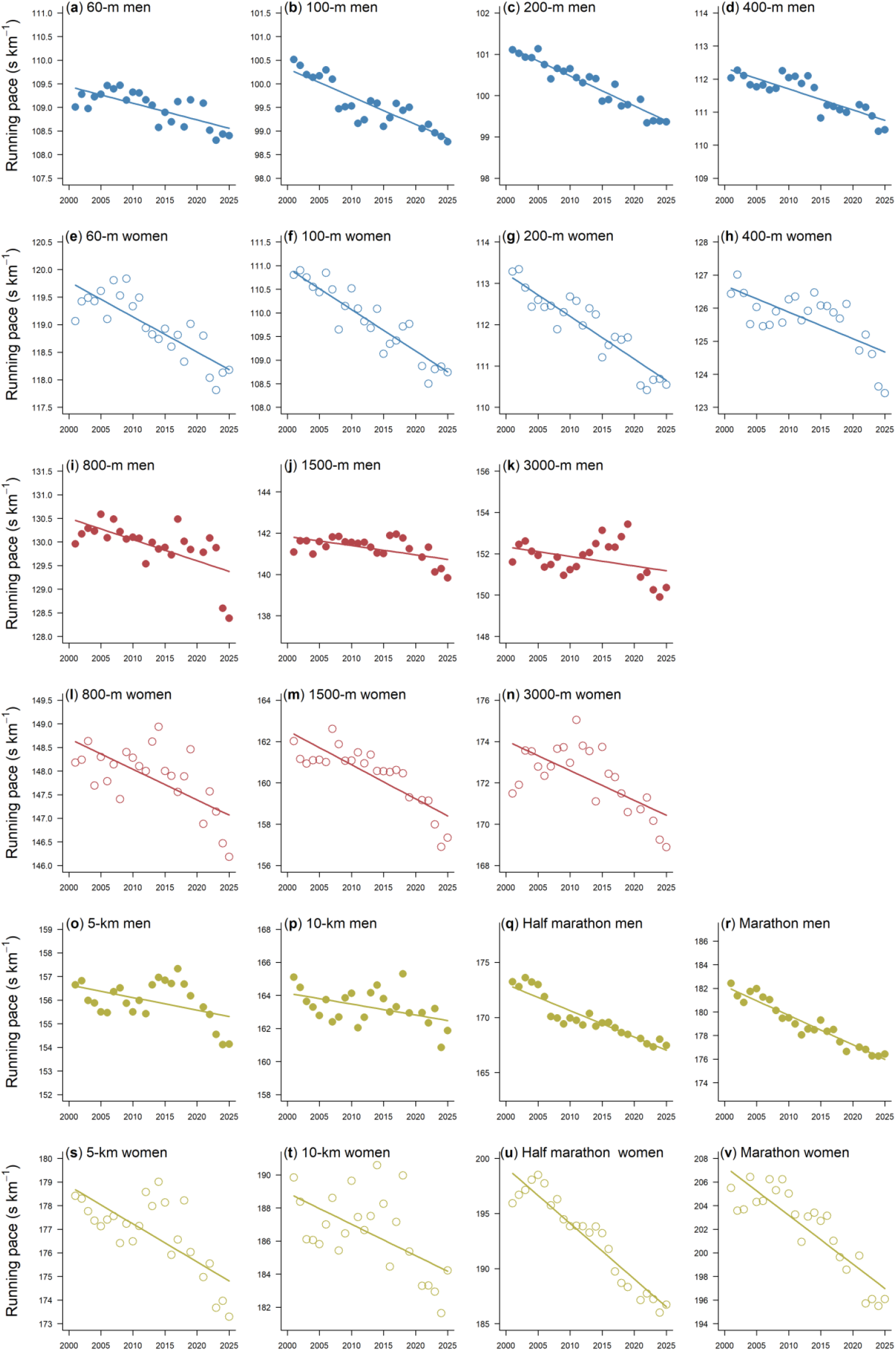
Progression in running pace of elite runners during the 21^st^ century. Changes in running pace between 2001 and 2025 for men (filled circles) and women (empty circles) across the 11 disciplines considered in this analysis. Each point corresponds to the mean running pace of the top 25 performing athletes within a given year. Running events are colour-coded into sprint (blue, top rows), middle-distance (red, middle rows) and long-distance races (yellow, bottom rows). Lines show the fit of the linear mixed effects models presented in Table S1. For each distance event, the range of the y-axis shown in the men’s (top) and women’s panels (bottom) is the same, making the regression slopes directly comparable between each pair of panels.

**Fig. 3:**
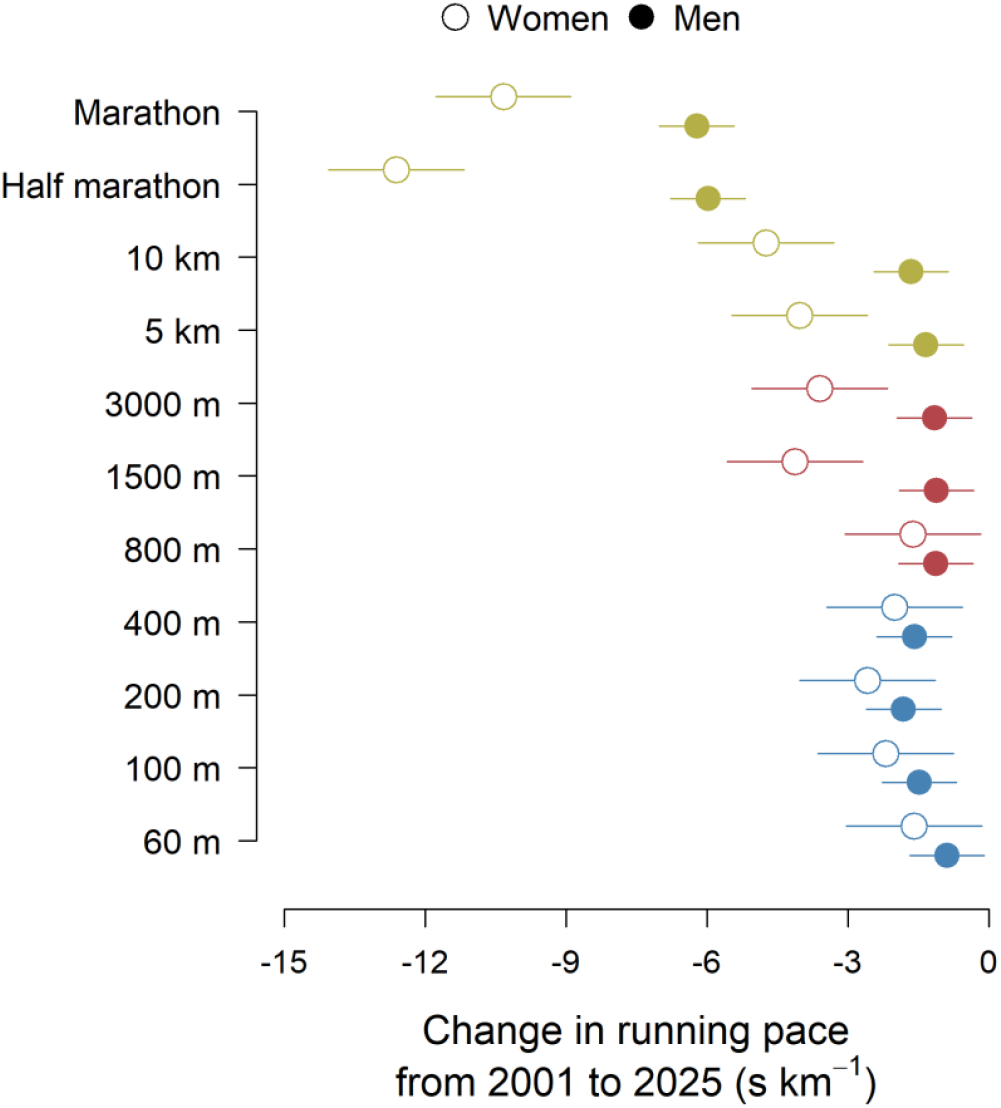
Increased running performance among elite athletes during the 21^st^ century. Change in running pace for female (open circles) and male (filled circles) athletes across the 11 events considered in this analysis for the period between 2001 and 2025. The more negative the values, the greater the increase in pace over the past 25 years. Values are estimates from linear mixed effects models, with error bars denoting 95% confidence intervals. Running events are colour-coded into sprint (blue), middle-distance (red) and long-distance races (yellow). Data underpinning estimated changes in running pace between 2001 and 2025 are shown in Fig. 2.

## Discussion

Both men and women have continued to push the frontiers of athletic performance during the 21^st^ century (13). Many people will remember Usain Bolt obliterating the 100 m world record in 2009, or when Eliud Kipchoge ran a marathon in under 2 hours in 2019. But progress by elite female athletes over this period has simply been much greater (Figs 2–3). Since the turn of the century, women have substantially improved almost all middle and long-distance world records (2.8% improvement on average compared to 1.3% for men; Table S2), running below 14 minutes in the 5 km and 29 minutes in the 10 km for the first time. Women have also improved the marathon world record by nearly 11 minutes since 2000 (7.7% improvement), with Ruth Chepngetich’s time of 2:09:56 being fast enough to win the men’s Olympic marathon as recently as 2004 (and at almost every other Olympics before then).

As previous studies have shown, the performance gap between men and women varies considerably with running distance (Fig. 1 a–b), being narrowest in short sprint events (8, 14). This likely reflects advantages of having shorter limbs and faster leg turnover rates when accelerating at the start of a race (8, 15). A perfect example of this are the 2024 Olympic 100 m finals, during which Noah Lyles and Julien Alfred – who won their respective races – reached the 10 m mark in almost the exact same time (1.95 s and 1.96 s) but where then separated by nearly a full second at the end of the race (9.79 s and 10.72 s). By contrast, as running distance increases, the greater muscle mass and aerobic capacity of male athletes initially results in a widening performance gap (3–6). Differences in running pace between men and women eventually stabilise in events longer that 5 km, meaning that in relative terms the performance gap actually shrinks in the half marathon and marathon (Fig. 1a–b). This is consistent with women having certain advantages over men in endurance races that can bridge the performance gap in ultramarathon events (5, 16–18).

Yet far from being static (6, 9–11, 19), the performance gap between elite male and female athletes has continued to evolve considerably since the turn of the century (Fig, 1c–d). This trend has been predominantly driven by sizable improvements by women in middle- and long-distance events (Figs 2–3). There are several factors that could have plausibly contributed to this, ranging from improved training regimes to the use of performance enhancing drugs (although anabolic steroids – which do disproportionately benefit women – are predominantly used to enhance muscle mass for short-distance events requiring explosive force; (20). But one factor in particular would seem likely to have played an important role: the recent development of advanced running shoes (so called ‘super shoes’). These first became adopted for road races in 2016 after the launch of the Nike Vaporfly, and have subsequently also been developed for track events since 2021. There is strong evidence that super shoes improve running economy and biomechanics (21–23), with performance gains being particularly pronounced in longer distance events (12, 23– 25). Crucially, studies have also found that super shoes may benefit female athletes more than men (12, 22, 26) – potentially explaining the trends in the data.

However, if super shoes were driving the shrinking gap between elite male and female athletes, the data should show a breakpoint around 2016 for road races and 2021 for track events, after which we would expect to see marked improvements by both sexes, with those by women being most pronounced. Instead, performance improvements by both men and women pre-date the advent of super shoes, with running pace having largely progressed linearly since the turn of the century across a large majority of running disciplines, including long-distance road races (Fig. 2). This suggest that while supershoes have undoubtedly played an important role in driving increased running performances in recent years, their use does not explain why elite female runners have improved so much more than men. In this regard it is worth noting that previous studied that have inferred that women benefit most from the use of super shoes have not been based on controlled experiments where the same athletes were made to run with and without super shoes. Instead, these studies calculated performance gains from the use of super shoes by comparing race times before and after this technology was introduced (12, 22, 26). The issue with this approach is that if running performances were already on a steady improvement trajectory before super shoes came into use, then one might mistakenly attribute gains in performance by women to the use of this technology when they are in fact driven by something else entirely.

In the absence of clear evidence that the shirking gap between elite male and female athletes coincided with the advent of super shoes, another plausible explanation for this trend is that it reflects, at least in part, a legacy of the different opportunities that men and women have historically had to participate in sport – especially over longer endurance events. Supporting this hypothesis with data is challenging, as participation in sport is influenced by a complex interplay of socio-economic and cultural factors (e.g., societal norms, lack of role models, access to training infrastructure, monetary incentives). But there are nonetheless several lines of evidence that are consistent with this idea. While it is true that landmark legislation improving access to sport for women such as Title XI in the US date to the early 1970s, the effects of these policies inevitably take decades to manifest. For example, women only began competing in the 5 km, 10 km and marathon at the Olympics as recently as 1996, 1988 and 1984, respectively. Similarly, comprehensive surveys of sports participation such as the State of Running report (27) reveal that the proportion of female runners has increased steadily from the mid-1980s (when it was below 20%) to present day (with women achieving parity with men in terms of participation as recently as 2018). Moreover, in terms of monetary incentives, we know from surveys of athletes that women continue to systematically earn less than men in athletics and that most professional female athletes are unable to live off their sponsorship deals alone (28, 29).

This suggests that as women’s professional running continues to grow, we may well see the performance gap between men and women shift towards a new equilibrium that has more to do with biology than opportunity (6, 30). Regardless of whether women will one day outrun men, what the data do strongly suggest is that over the coming decades women will continue to redefine the boundaries of athletic performance. Currently Faith Kipyegon is bidding to become the first woman to run under 4 minutes for the mile, emulating what Sir Roger Bannister did in 1954 in what is surely the most iconic moment in modern running history to date. I would not bet against her.

## Acknowledgements

TJ was supported by a UK NERC Independent Research Fellowship (NE/S01537X/1) and through a UKRI Frontier Research grant (EP/Y003810/1).

