## Supporting Information for "The gap between elite female and male runners continues to shrink in the 21st century"

**Table S1**: Summary outputs of linear mixed effects models testing how the gender gap (expressed both as a % and a pace, in s km-1), as well as male and female running paces have changed over time between 2001 and 2025. Each model included calendar year as a fixed effect, as well as a random slope and intercept term that allowed the relationship to vary among the 11 running events (coded as '*year | event*' in the *lme4* package in R). To aid model convergence and interpretation, prior to model fitting year was rescaled so that the timeseries start (2001) was set to 0. Consequently, the intercept term of the model can be interpreted as the estimated gender gap or running pace in 2001, while the slope corresponds to the annual change in gender gap or running pace. *P* values for fixed effects terms were calculated using the *lmerTest* package, which implements Satterthwaite's approximation for degrees of freedom. For scatterplots of data and model fits, see Fig. 2 in the main text.

| **Model** | **Fixed effect** | **Parameter value (±95% CI)** | ***P* value** |
|---|---|---|---|
| Gender gap (pace) ~ year + (year \| event) | Intercept | -18.65 (±3.49) | <0.001 |
| | Slope | 0.091 (±0.048) | 0.004 |
| Gender gap (%) ~ year + (year \| event) | Intercept | -13.20 (±1.09) | <0.001 |
| | Slope | 0.053 (±0.024) | 0.002 |
| Men's pace ~ year + (year \| event) | Intercept | 138.5 (±17.4) | <0.001 |
| | Slope | -0.089 (±0.048) | 0.005 |
| Women's pace ~ year + (year \| event) | Intercept | 157.1 (±20.8) | <0.001 |
| | Slope | -0.180 (±0.090) | 0.003 |

**Table S2**: Progression of men's and women's world records between 2000 and 2025 across middle and long-distance events ranging from 800 m to the marathon. Progressions are reported both in seconds (s) and as proportional improvements (%). Data obtained from the World Athletics website (https://worldathletics.org).

| **Event** | **World record progression between 2000 and 2025 (seconds and %)** | |
|---|---|---|
| | **Men** | **Women** |
| 800 m | 0.2 s (0.2 %) | 0 s (0 %) |
| 1500 m | 0 s (0 %) | 1.8 s (0.8 %) |
| Mile | 0 s (0 %) | 6.1 s (2.4 %) |
| 3000 m | 3.1 s (0.7 %) | 0 s (0 %) |
| 5 km | 4 s (0.5 %) | 30 s (3.5 %) |
| 10 km | 11.8 s (0.7 %) | 37.6 s (2.1 %) |
| Half-marathon | 144 s (4.1 %) | 232 s (5.8 %) |
| Marathon | 307 s (4.1 %) | 647 s (7.7 %) |

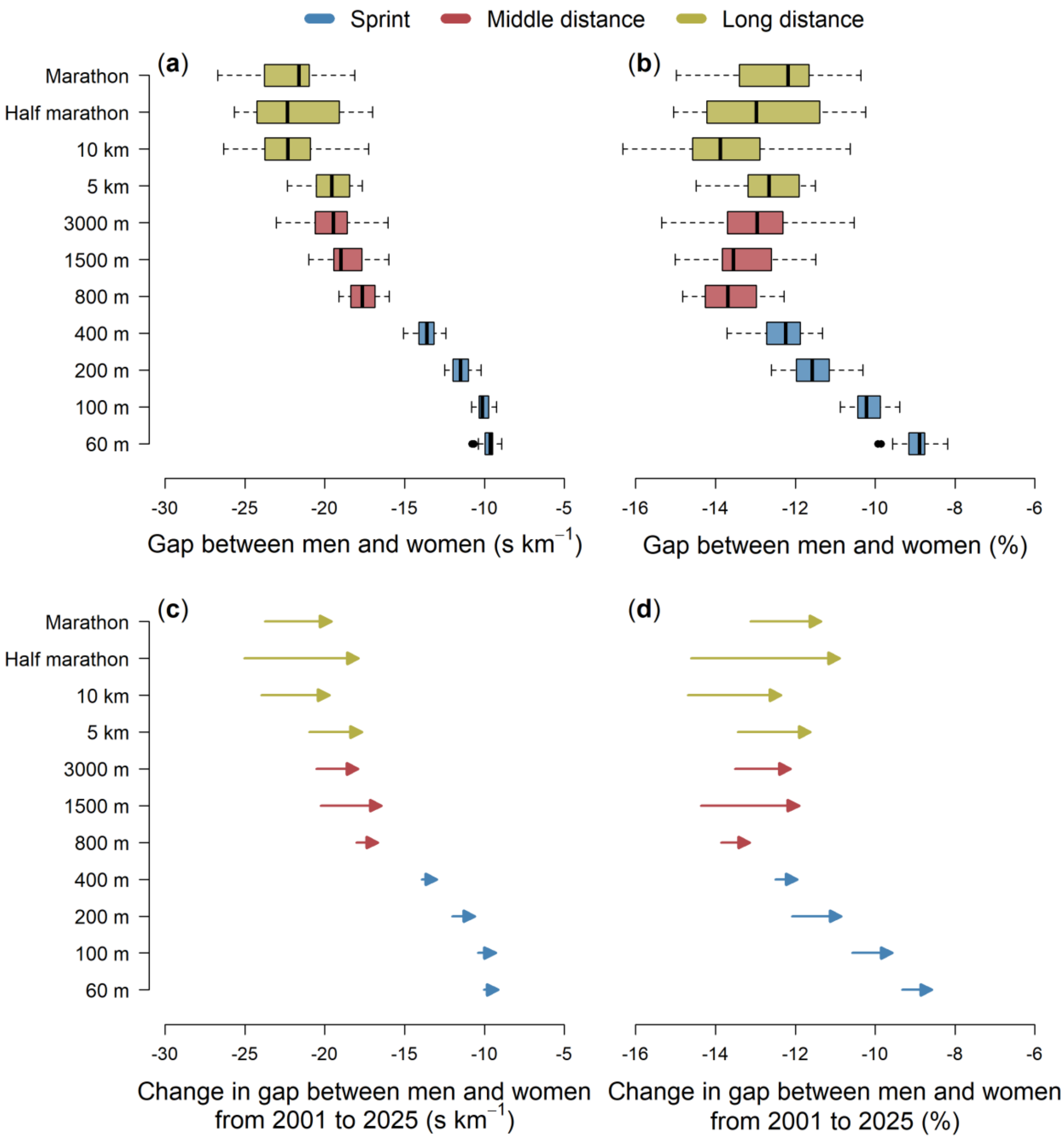


**Fig. S1**: This figure is analogous to Fig. 1 in the main text, but is calculated using data for the top 10 performing athletes across each discipline (instead of the top 25). Top panels depict variation in the gap – expressed as both a difference in pace between men and women (**a**) and as a % (**b**) – across the 11 events considered in this analysis for the period between 2001 and 2025. The more negative the values, the greater the gap between male and female runners. Bottom panels (**c–d**) show changes in the gap between 2001 (start of the arrow) and 2025 (arrow head) for each running event, as estimated from linear mixed effects models. Arrows pointing to the right indicate a shrinking gap over time, with the length of the arrow reflecting the magnitude of this change. Running events are colour-coded into sprint (blue), middle-distance (red) and long-distance races (yellow).

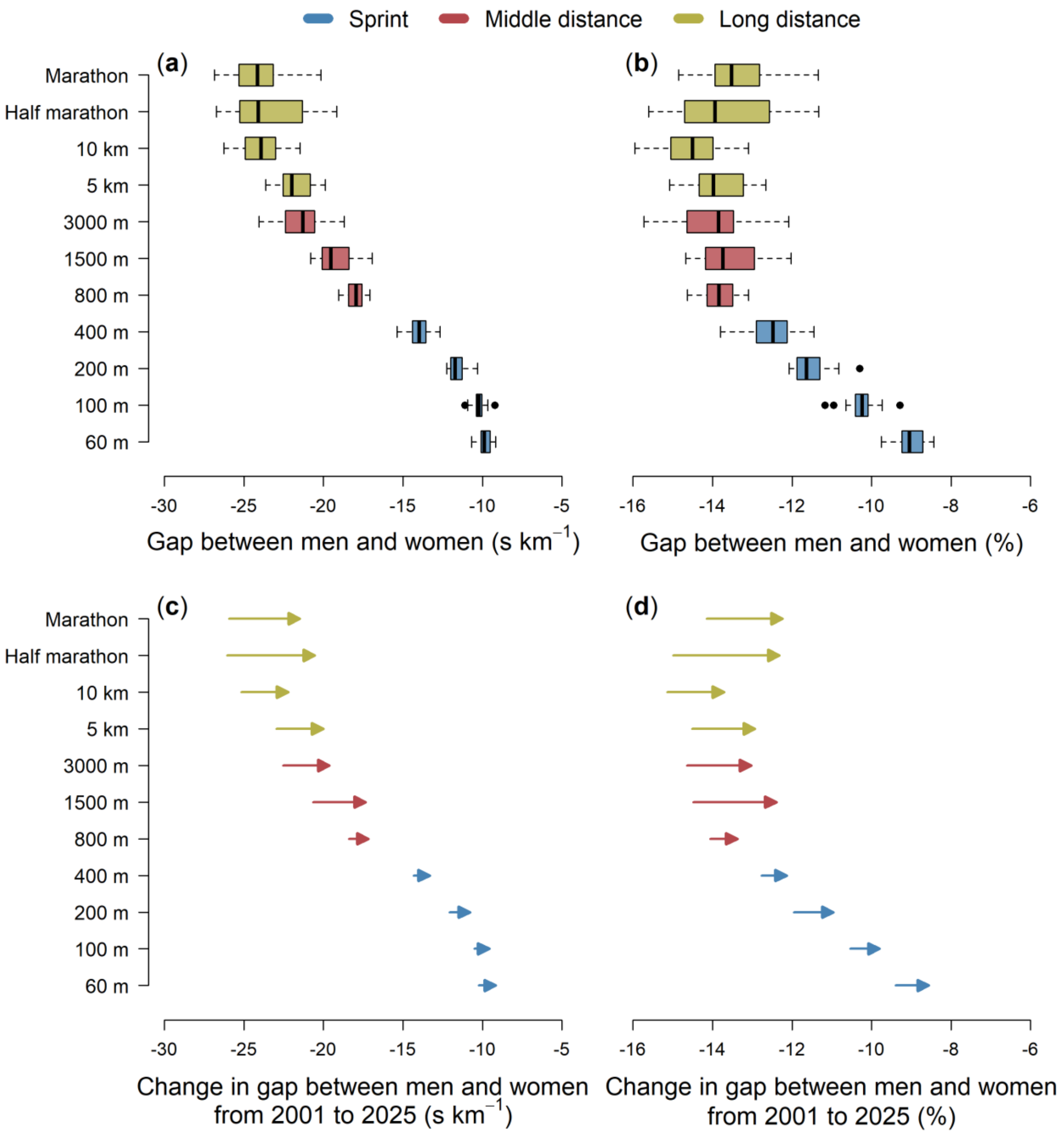


**Fig. S2**: This figure is analogous to Fig. 1 in the main text, but is calculated using data for the top 50 performing athletes across each discipline (instead of the top 25). Top panels depict variation in the gap – expressed as both a difference in pace between men and women (**a**) and as a % (**b**) – across the 11 events considered in this analysis for the period between 2001 and 2025. The more negative the values, the greater the gap between male and female runners. Bottom panels (**c–d**) show changes in the gap between 2001 (start of the arrow) and 2025 (arrow head) for each running event, as estimated from linear mixed effects models. Arrows pointing to the right indicate a shrinking gap over time, with the length of the arrow reflecting the magnitude of this change. Running events are colour-coded into sprint (blue), middle-distance (red) and long-distance races (yellow).
